# Sensitive, reliable, and robust circRNA detection from RNA-seq with CirComPara2

**DOI:** 10.1101/2021.02.18.431705

**Authors:** Enrico Gaffo, Alessia Buratin, Anna Dal Molin, Stefania Bortoluzzi

## Abstract

Circular RNAs (circRNAs) are a large class of covalently closed RNA molecules that originate by a process called back-splicing. CircRNAs are emerging as functional RNAs involved in the regulation of biological processes as well as in disease and cancer mechanisms. Current computational methods for circRNA identification from RNA-seq experiments are characterised by low discovery rates and performance dependent on the analysed data set. We developed a new automated computational pipeline, CirComPara2 (https://github.com/egaffo/CirComPara2), that consistently achieves high recall rates without losing precision by combining multiple circRNA detection methods. In our benchmark analysis, CirComPara2 outperformed state-of-the-art circRNA discovery tools and proved to be a reliable and robust method for comprehensive transcriptomics characterisation.

## Introduction

Recent research uncovered that eukaryotic transcriptomes comprise thousands of stable circular RNAs (circRNAs) that originate by a process called back-splicing, where the transcript 3’ and 5’ ends are covalently joined (Xiao et al. 2020). Rather than being transcriptional byproducts, circRNA molecules exert critical functions in cell biology through different mechanisms (Bonizzato et al. 2016). By interacting with microRNAs, circRNAs can regulate gene expression and govern important oncogenic axes (Hansen et al. 2013); moreover, similarly to long non-coding RNAs, they can control diverse cellular processes by decoying RNA-binding proteins and scaffolding molecular complexes (Du et al. 2016). CircRNAs can also function as templates for translation to encode functional peptides (Xiao et al. 2020)(Wu et al. 2021b) and regulate the transcription of their parental gene (Li et al. 2015). Nowadays, circRNAs are considered key players that can impact all the cancer hallmarks (Hanniford et al. 2020; Slack and Chinnaiyan 2019). The discovery of circRNA regulatory roles and their potential as biomarkers given by higher stability compared to linear RNAs (Rajappa et al. 2020) have actuated the integration of circRNA investigation in conventional transcriptomics, especially in cancer research and studies of pathological conditions (Santer et al. 2019; Hua et al. 2019), including viral infections (Awan et al. 2021).

Studies of circRNAs rapidly increased in pace thanks to the development of bioinformatics tools that identify the sequences spanning circRNA back-splice junctions from total RNA-seq data. To date, several methods for circRNA identification have been developed (Chen et al. 2020). Still, none outperforms the others since they all provide either highly sensitive or highly precise predictions and highly variable performance across benchmark data sets (Chen et al. 2015; Hansen et al. 2016; Zeng et al. 2017). Interestingly, Hansen (Hansen 2018) observed that circRNA detection methods largely agreed on true predictions, whereas circRNAs identified by single methods were enriched in false-positive guesses (FPs) and suggested selecting circRNAs commonly predicted by two or more methods to obtain dependable results.

We formerly implemented CirComPara (Gaffo et al. 2017), an automated computational pipeline combining four circRNA detection methods, including CIRCexplorer (Zhang et al. 2014), CIRI2 (Gao et al. 2018), Findcirc (Memczak et al. 2013) and Segemehl (Hoffmann et al. 2014). CirComPara controlled the FP number by considering only the circRNAs commonly detected by two or more methods.

In CirComPara2, we have considerably improved our tool by: (i) including five additional circRNA detection methods, (ii) updating the software of the already integrated tools, (iii) implementing a more accurate counting of the back-spliced reads, (iv) increasing the analysis pipeline flexibility, and (v) including the procedure to calculate the linear expression related to the circRNAs.

In this work, we first show that nine widely used circRNA detection tools could miss circRNAs of interest. Then, we confirm that CirComPara2 correctly reports circRNAs overlooked by other methods and achieves significantly higher sensitivity with no loss of precision. Finally, assessment on simulated data and 142 public RNA-seq samples demonstrated the consistent higher performance of CirComPara2 compared with state-of-the-art methods.

## Results

### CircRNA detection methods could miss abundant circRNAs

We simulated RNA-seq expression data of 5,680 circRNAs from the whole human genome (“simulated data set”; see Methods) to evaluate the characteristics of circRNA detection methods’ false-negative predictions (FNs), i.e., true circRNAs not identified as such. We applied nine widely used computational pipelines for circRNA discovery including circRNA_finder (Westholm et al. 2014), CIRI2 (Gao et al. 2018), DCC (Cheng et al. 2016), Findcirc (Memczak et al. 2013), Segemehl (Hoffmann et al. 2014), and CIRCexplorer2 (Zhang et al. 2016). CIRCexplorer2 was coupled to any of BWA (Li 2013) (C2BW), Segemehl (C2SE), STAR (Dobin et al. 2013) (C2ST) and TopHat-Fusion (Kim and Salzberg 2011) (C2TH) aligners, thus composing four different pipelines.

On average, 49% of FNs detected by each method showed higher expression than the overall circRNA median expression (Figure 1a), suggesting that nearly half of the missed circRNAs had a considerable expression level no matter which method was applied. Besides, the expression distribution of the FNs was similar to the correctly identified circRNAs, i.e., the true-positive findings (TPs), whereas the false-positive (FP) expression was generally low.

**Figure 1.**
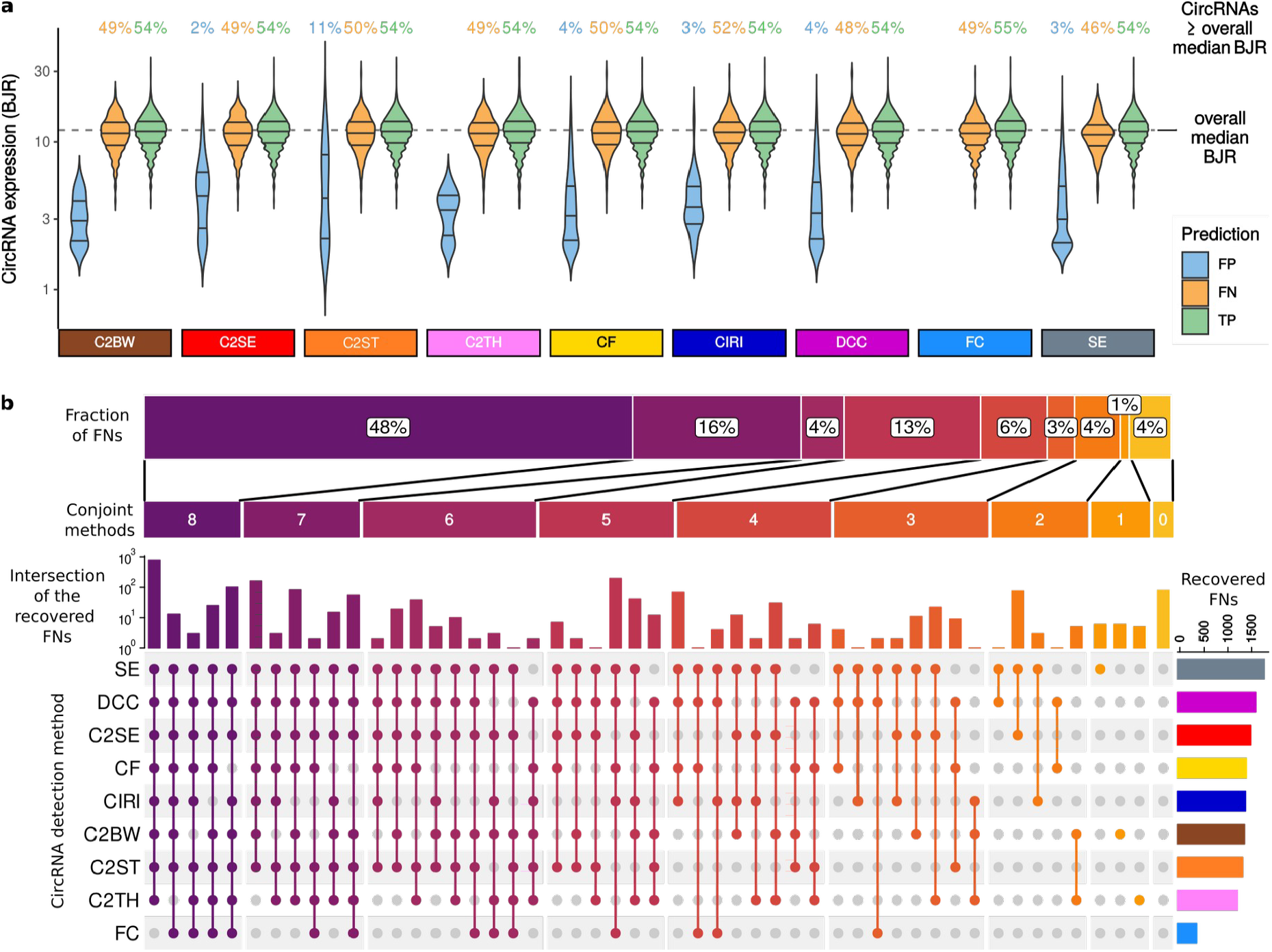
CircRNAs overlooked by common detection methods and the CirComPara2 approach. **a**, The expression level of predicted circRNAs. BJR: back-splice junction read counts; C2BW: CIRCexplorer2 on BWA; C2SE: CIRCexplorer2 on Segemehl; C2ST: CIRCexplorer2 on STAR; C2TH: CIRCexplorer2 on TopHat-Fusion; CF: circRNA_finder; CIRI: CIRI2; FC: Findcirc; SE: Segemehl; FP: false-positive; FN: false negative; TP: true positive. **b**, Number of methods detecting circRNAs not detected by other methods and the number of the FNs detected. Colour refers to the number of methods conjointly detecting circRNAs missed by other tools. The vertical bars show the number of FN circRNAs detected by the methods indicated in the coloured dots below the bars. The bars denote disjoint circRNA sets. Grey dots indicate the methods failing to detect the circRNAs considered in the bar chart on the top. The horizontal bars on the right represent the overall number of FN circRNAs detected by the methods. The horizontal bar shows the percentage of detected FNs by grouping method combinations according to the number of combined methods.

### Multiple methods compensate each other, and method combinations increase the detection sensitivity

We further examined the 1,945 circRNAs undetected by one or more tools, referred to as the “FN set” from now on, by counting how many circRNAs in the FN set each method could detect.

Interestingly, only 4% circRNAs of the FN set were undetected by all methods (Figure 1b), whereas 96% could be identified at least by one among the nine tools. Specifically, 1% FNs were detected individually by Segemehl, C2BW, and C2TH; and 95% were commonly identified by various combinations of two or more tools. Almost half the FNs (48%) were conjointly detected by eight out of nine methods, with Segemehl, DCC and C2SE providing the most inclusive predictions. Instead, Findcirc showed the least number of recovered FNs. However, no method entirely covered the predictions of Findcirc, indicating some specificity of its algorithm.

Overall, this analysis suggested that algorithms with possibly different and complementary features can compensate each other and improve the detection rate if applied together.

### Workflow and features of CirComPara2

Following the observation reported in the previous paragraph, we enhanced CirComPara by including more circRNA detection methods to improve its sensitivity. Moreover, we introduced new features that made CirComPara2 more flexible, computationally efficient, and resilient.

CirComPara2 implements a fully automated computational pipeline for circRNA detection, quantification, annotation, and integration with linear gene expression data (Figure 2a). Several parameters are available to customise the analysis workflow and the integrated methods. The minimal input consists of the RNA-seq reads in FASTQ format and a reference genome in FASTA format. Optionally, the user can also provide the gene annotation in GTF format. The software will then build the genome indexes for each read aligner and perform the necessary file format conversions. Previously computed indexes can be reused as input to save computing time.

**Figure 2.**
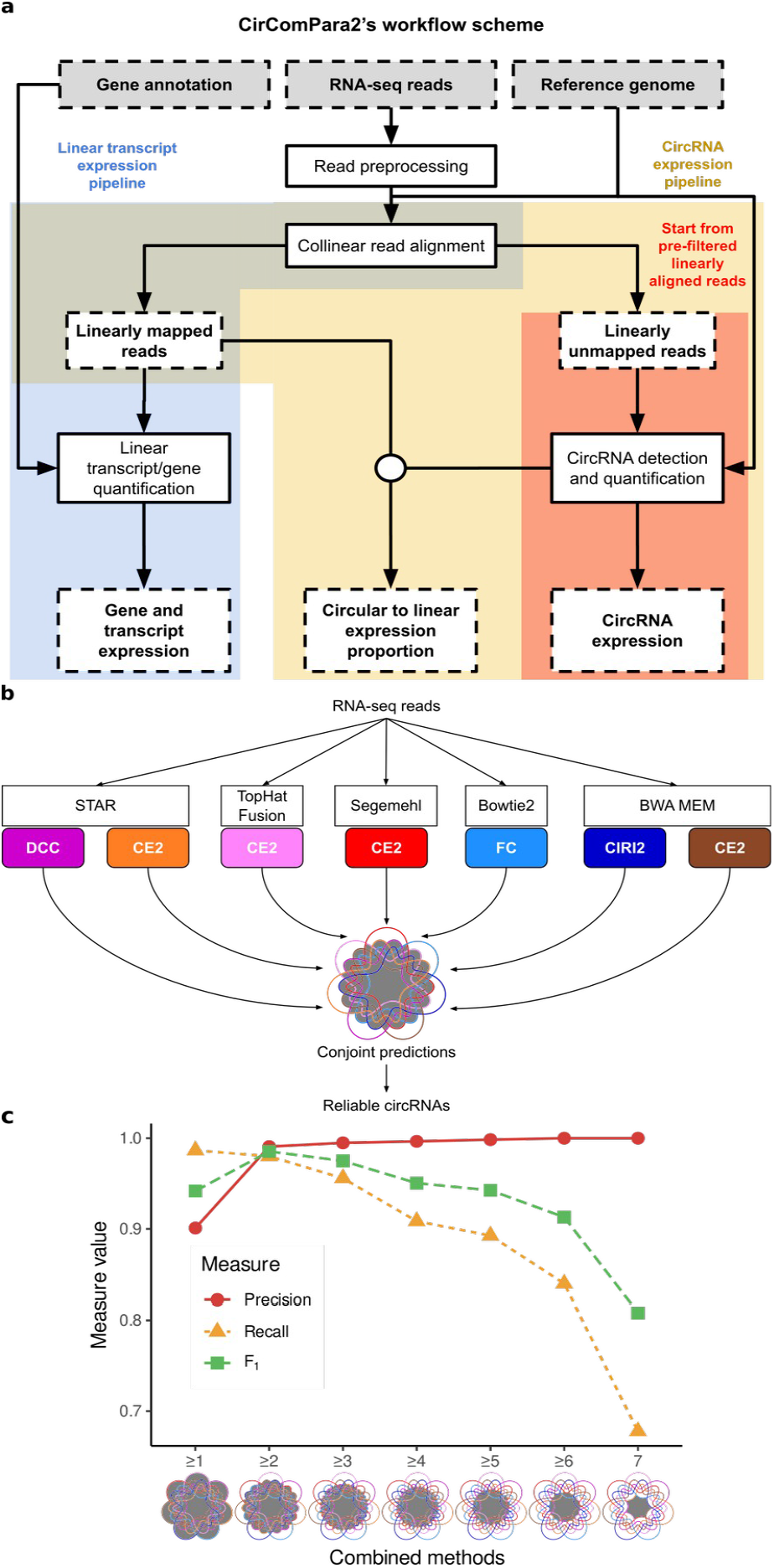
The CirComPara2 workflow and approach. **a**, Boxes with dashed contour indicate input and output data; boxes with solid-line contour indicate computing tasks; the circular connector indicates merging data. Background colours highlight the different pipeline branches (linear transcript analysis in blue, full circRNA analysis in yellow, and strict circRNA analysis in red). **b**, Detail of the CirComPara2 strategy (“CircRNA detection and quantification” box in **a**) showing the embedded circRNA detection methods (coloured rounded corner boxes) with the respective chimeric read aligners (white boxes). The central Venn diagram represents the optimised combination of method prediction intersections implemented by CirComPara2 with circRNAs conjointly detected by two or more methods (grey filled intersections) retained as default. Method abbreviations as in Figure 1. **c**, Precision (red), recall (yellow) and F_1_-score (green) for different numbers of methods considered by CirComPara2: ≥1 indicates the union of all single methods predictions, ≥2 indicates any pair of methods, and so on, up to 7, which indicates the conjoint predictions of all the methods considered. The grey filled parts in Venn diagrams show the intersections considered by the conjoint method combinations.

The CirComPara2 workflow comprises an optional pre-processing of the input raw reads by Trimmomatic (Bolger et al. 2014) to trim or discard low-quality reads. Statistics of the preprocessing steps are produced with the FastQC tool (Andrews 2010). Next, the reads are aligned collinearly to the reference genome using HISAT2 (Kim et al. 2019) to (i) identify the reads that are later used for linear transcript analysis (Figure 2a) and (ii) extract the reads not collinearly aligned, which are used to detect back-splices. The linear gene and transcript expression analysis is performed with StringTie (Pertea et al. 2015) and produces files that can be easily imported into packages for downstream expression analysis such as *tximport* (Soneson et al. 2015) and *tximeta* (Love et al. 2020). The circRNA analysis aligns the collinearly unmapped reads independently with five methods allowing chimeric alignments, namely Bowtie2 (Langmead and Salzberg 2012), BWA-MEM, Segemehl, STAR and TopHat-Fusion (Figure 2b). The chimeric aligner outputs are subsequently parsed by six circRNA detection tools, which compose the nine different circRNA detection sub-pipelines presented in the previous paragraph. Of note, the computationally expensive chimeric alignment step is performed only once per aligner and reused by multiple circRNA detection tools, boosting CirComPara2 efficiency. For instance, the same alignments from STAR are passed to CIRCexplorer2, circRNA_finder, and DCC (Figure 2b). The outputs of the various tools are automatically handled, converting them into a standard format to compare the predicted back-splices. Moreover, the identifiers of the back-spliced reads are collected while keeping track of the predicting method to obtain non-redundant read counts for each circRNA. Finally, the linear expression of circRNA host genes is evaluated by counting the reads collinearly mapped at each back-splice junction using bedtools (Quinlan and Hall 2010), GNU parallel (Tange 2020), and custom scripts.

CirComPara2 has a modular and highly parallelised implementation that makes it computationally efficient and resilient. By using custom parameters, CirComPara2 allows skipping computation tasks that are not of interest to the user. For instance, the user can select to run only the pipeline branch computing the linear or the circular transcript analysis, the collinear alignments (for instance, if they were previously computed), or both the collinear alignment and the linear transcript pipeline branch, therefore performing only the circRNA detection from pre-filtered reads. Plus, the Scons (www.scons.org) engine is leveraged to run independent tasks in parallel, resume an interrupted analysis by performing only uncompleted tasks, and, if the user changed some parameters from a previous run, compute only the tasks dependent on the modified parameters.

CirComPara2 is available as standalone software (https://github.com/egaffo/circompara2) and Docker image (https://hub.docker.com/r/egaffo/circompara2), which facilitates installation, portability, and reproducibility of the analysis.

### Optimal method combination for circRNA detection

Concurrently to the improvement of CirComPara2 sensitivity, we wanted to control the number of the introduced FPs to preserve a high precision. As observed in a previous study (Hansen 2018), the combination of specific methods does not ameliorate the discovery of true circRNAs. Accordingly, our method combination assessment on the simulated data showed that combining circRNA_finder or Segemehl with other methods contributed to increasing the FP number (Supplementary Figure S1). For this reason, circRNA_finder and Segemehl were excluded in the CirComPara2 default method combination (Figure 2b); nevertheless, these two methods can be included if enabled by the user.

Next, we fine-tuned the number of conjoint methods considering the amount of recovered FNs against the introduced FPs to optimise the method combination strategy. As expected, considering the predictions from all methods resulted in the highest recall (0.99) and the lowest precision (0.90) among the combination strategies (Figure 2c). Further, excluding predictions from single methods, i.e., selecting only the circRNAs commonly detected by two or more tools, showed a slightly reduced recall (0.98) with a substantially increased precision (0.99). This large precision gain indicated that most of the FPs were predicted by single methods. Further increasing the number of conjoint methods (i.e., from three-or-more to all-seven methods conjoint) lead to a considerable decrease of recall (0.96 to 0.68) with only a modest increase of precision (0.99 to 1.00).

To evenly weigh recall and precision in ranking the method combination strategies, we calculated the F_1_-score for each combination. The best trade-off between recall and precision, indicated by the highest F_1_-score (0.99), was obtained with the two-or-more method strategy (Figure 2c).

### CirComPara2 outperforms other methods in simulated data

We next set CirComPara2 with the two-or-more method strategy and compared it with the single methods on the simulated data. CirComPara2 obtained the highest F_1_-score (0.99) by achieving the highest recall (0.98) while holding a precision comparable to or higher than the other algorithms (0.99 versus 0.92–1.00; Figure 3a), confirming that CirComPara2 rectified the true circRNAs missed by the other methods.

**Figure 3.**
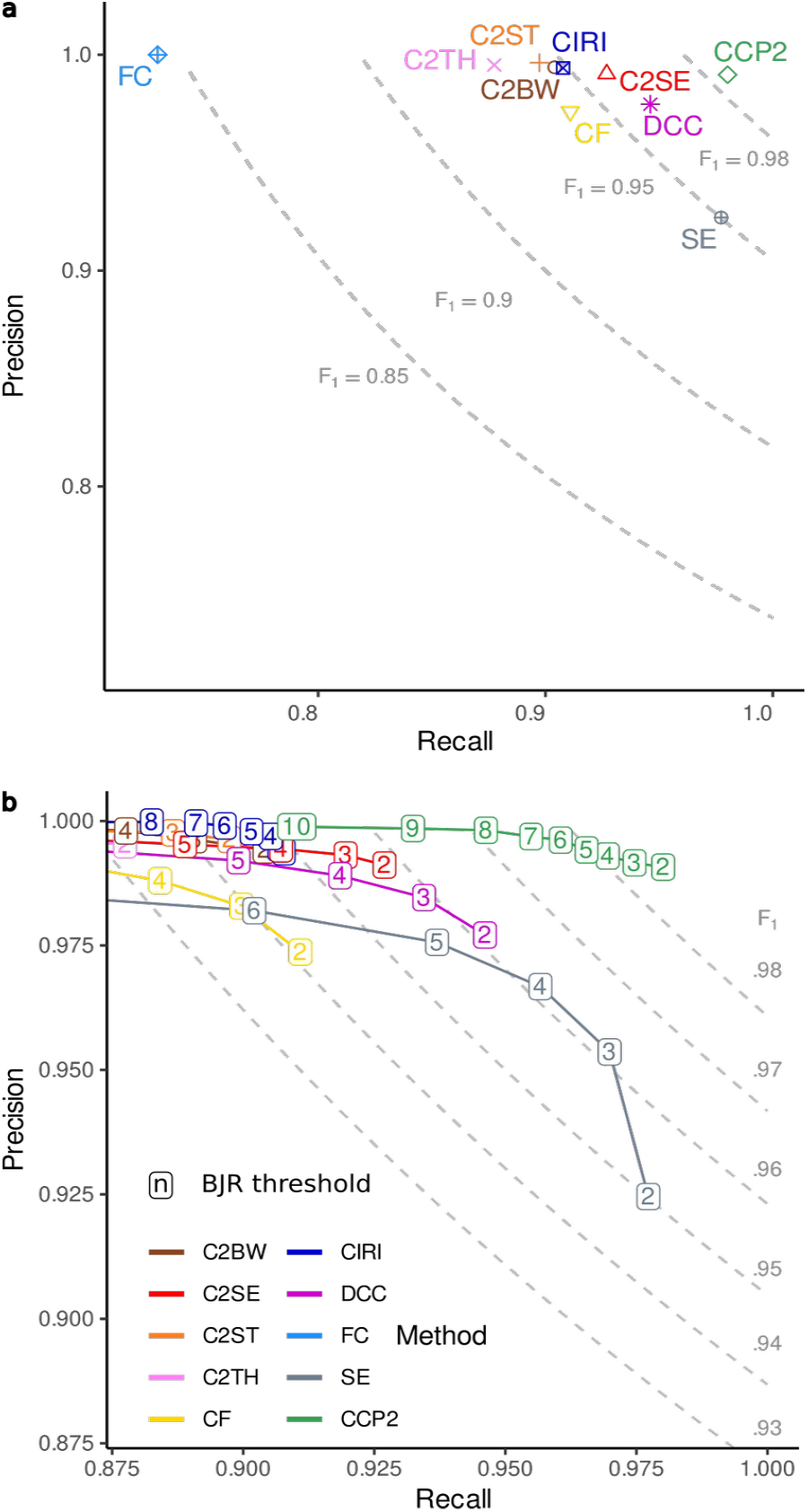
Performance of circRNA detection methods on the simulated data set. **a**, Precision and recall of the circRNA detection methods (labels as in Figure 1), including CirComPara2 (CCP2; green dot); the dashed-line curves display plot areas for 0.85, 0.90, 0.95 and 0.98 F_1_-scores. **b**, CircRNA detection methods’ performance upon application of filters from 2 to 10 minimum circRNA raw expression estimates (BJR: back-splice junction read counts); dashed-line curves delimit F_1_-scores plot areas from 0.92 to 0.98 with 0.01 increase steps; method colours as in (a).

To assess the extent of the annotation-guided method contribution to CirComPara2 predictions, we performed the analysis also with a pruned gene annotation input to the algorithms (see Methods). As expected, the CIRCexplorer2 pipelines showed a dramatic reduction (up to −0.14) of the recall and F_1_-scores (Supplementary Figure S2). Instead, CirComPara2 maintained the highest F_1_-score (0.98), suggesting that it can be efficient when applied to RNA-seq data of organisms with incomplete or poor genome annotation by leveraging the embedded annotation-independent tools.

A typical circRNA expression analysis usually involves post-detection data cleaning to remove background noise signals or mapping errors and poorly expressed circRNAs of little interest (Chen et al. 2016). Consequently, the circRNAs with small back-splice junction read counts (BJRs) are routinely filtered out. As shown in Figure 1a, FPs generally have small BJRs, suggesting that more reliable circRNAs can be retained by simply filtering according to expression abundance. We applied this procedure to our data, progressively filtering out circRNAs with less than two up to 10 BJRs. We stress that removing circRNAs with ≤10 BJRs was an extreme filter since it was close to the overall median BJR (Figure 1a). As expected, by increasing the minimum-BJR filter threshold, all methods scored higher precision but with a corresponding reduced recall, indicating that true circRNAs were discarded as well (Supplementary Figure S3). Nevertheless, CirComPara2 maintained the highest recall compared to other methods at equal precision (Figure 2b; Supplementary Table S1), suggesting that the circRNAs recovered by CirComPara2 were valid findings with a considerable abundance that may represent relevant circRNAs in actual experiments.

### Benchmark real data sets

The gold standard for evaluating circRNA detection methods on real RNA-seq data is to compare ribosomal RNA-depleted (ribo^-^) with circRNA-enriched sequencing libraries. The most used technique is the additional treatment of the ribo^-^ library with RNase R to exploit the exonuclease degradation resistance of circRNAs derived by their lack of a single-stranded 3′ end (Hansen et al. 2016; Hansen 2018; Gao et al. 2018; Zeng et al. 2017; Szabo and Salzman 2016). Accordingly, we collected a total of 142 public real RNA-seq data samples for which sample-matched ribo^-^ and RNase R-treated libraries were available (Table 1). This validation data set comprised samples of human cell lines and various human, Rhesus macaque, and mice tissues from six independent studies.

**Table 1.**
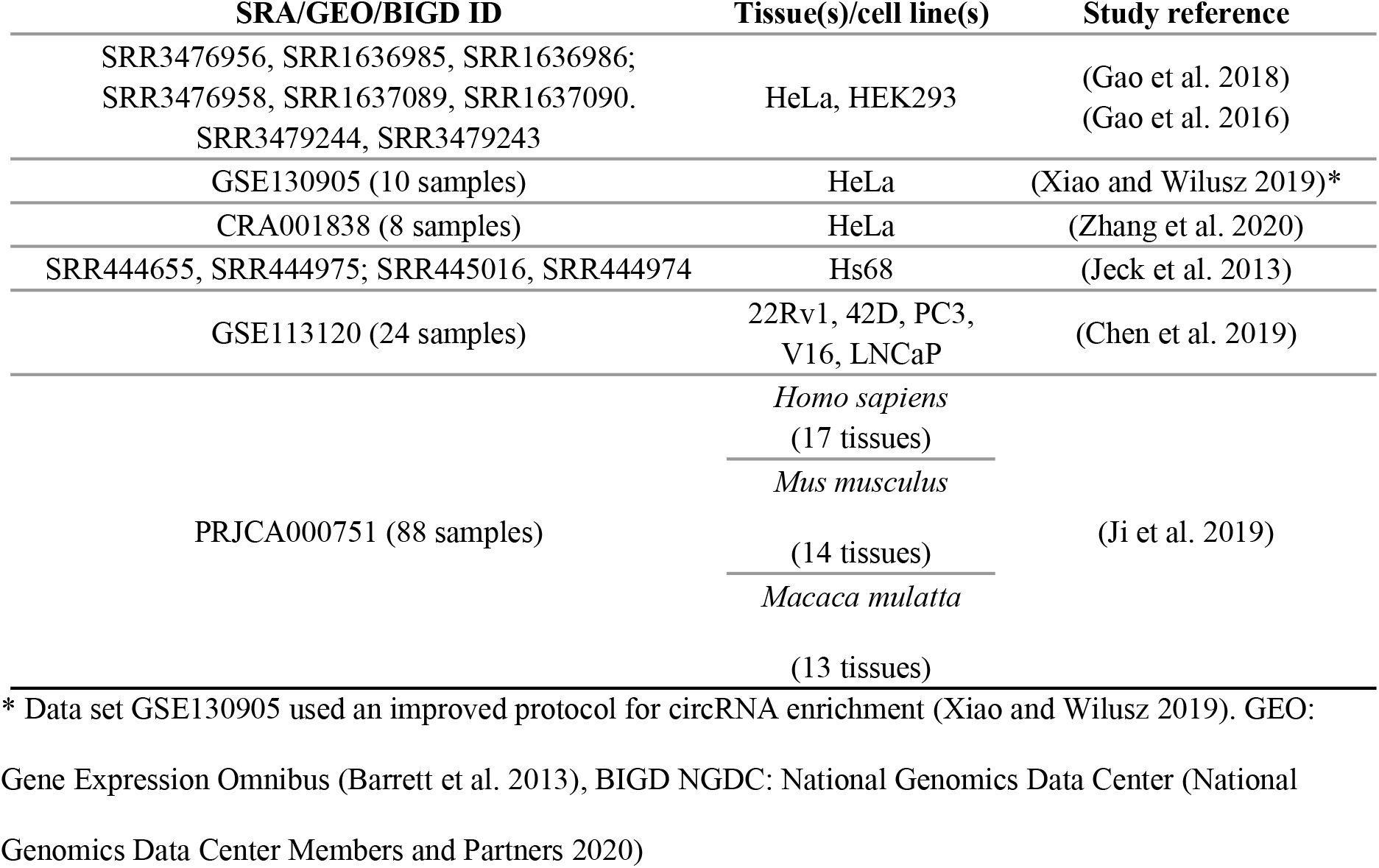
Real data sets of rRNA- and rRNA-/RNase R treated matched pairs of samples used to benchmark the circRNA detection methods. Tissues and cell lines are from humans unless specified.

The matched RNase R-treated samples were used as a control to define the TP circRNAs (see Methods). Assuming that the RNase R treatment would deplete the linear more than the circular transcripts, we considered TPs the circRNAs having circular-to-linear expression proportion (CLP) higher or equal in the RNase R-treated compared to the ribo^-^ matched samples according to at least one method. In this way, we accounted for different sequencing library depths between the matched samples. Moreover, each circRNA host-gene linear expression was estimated commonly to all the detection tools and independently of the circRNA abundance estimated by each method, allowing to remove possible advantages given by specific method quantification approaches.

### CirComPara2 performs consistently better than single methods

The performance of the methods on the real RNA-seq data agreed with the analysis on simulated data and previous benchmark study results (Zeng et al. 2017; Hansen 2018). Segemehl showed high recall (median 0.75; Figure 4a) but the lowest precision (median 0.92; Figure 4b), whereas C2BW, as expected from annotation-based algorithms, showed the most reliable predictions (median precision 0.97; Figure 4b). Similar to the evaluation on the simulated data set, we computed the F_1_-scores on the real data set. According to F_1_-score medians, CirComPara2 had the highest (0.91) and significantly different value (q < 0.001; Figure 4c), and substantially outperformed the next best method, Segemehl (median F_1_-score 0.82), with a 0.09 F_1_-score difference (Supplementary Table S2). The highest F_1_-score, achieved by CirComPara2, resulted from a significantly larger recall (median 0.86; q < 0.001; Figure 4a) and a negligible loss of precision compared to the other methods (0.01 median reduction to the most precise method; Figure 4b). Moreover, CirComPara2 had the narrowest interquartile range of F_1_-scores across the real-data sets (0.11; Figure 4c; Supplementary Table S2), proving that it is robust and almost unaffected by the experimental scenario. In light of the more challenging nature of real compared to simulated data (Szabo and Salzman 2016; Chen et al. 2015), these results further confirmed the advantage of CirComPara2 over the methods here assessed.

**Figure 4.**
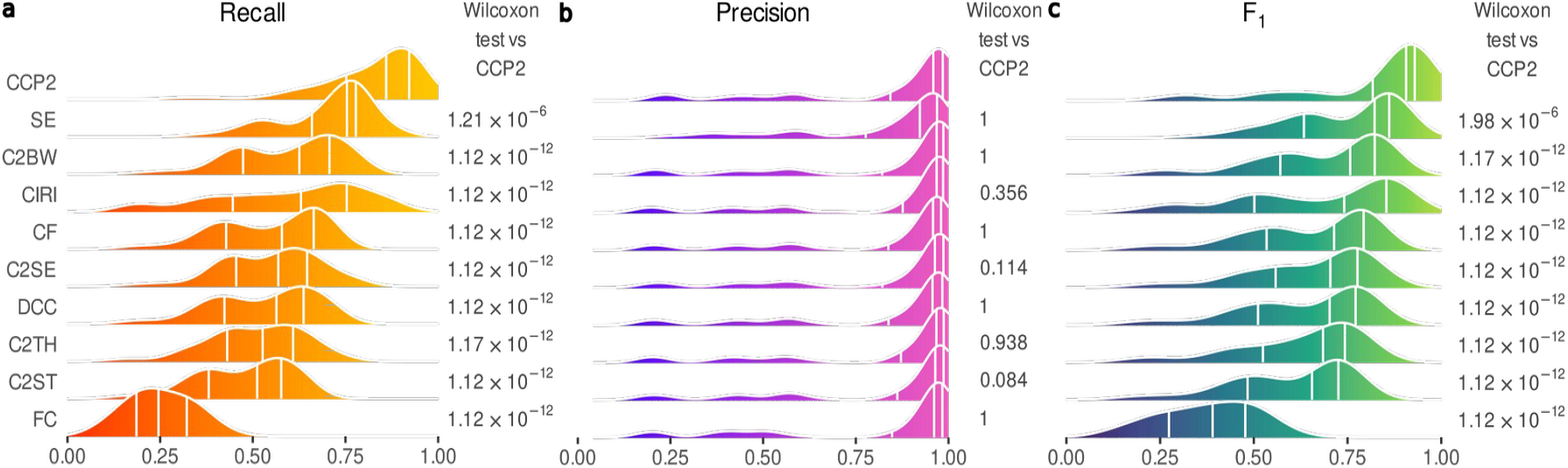
Performance of circRNA detection methods on the real RNA-seq data set. Results of the analysis on the 71 pairs of samples of the real RNA-seq data set. Density plot of (**c**) recall, (**d**) precision and (**e**) F_1_-score; distribution quartiles are indicated by white vertical lines. On the right-hand side of each plot, the Bonferroni adjusted p-values of Wilcoxon paired test contrasting each method with CirComPara2. Methods’ abbreviations as in Figure 1; CCP2: CirComPara2.

## Discussion

In the early days after circRNA discovery, bioinformaticians put significant effort into developing circRNA detection methods with highly precise predictions. To this aim, the current circRNA detection algorithms apply filtering procedures to remove FPs, but that may also reject true circRNAs (Zeng et al. 2017; Chen et al. 2015; Hansen 2018; Hansen et al. 2016), increasing the number of FNs and the risk of overlooking circRNAs of interest, as suggested by our analysis. Moreover, the frequent practice of filtering out low count circRNAs to improve precision may result in the loss of differentially expressed elements in comparative studies, as demonstrated by systematic studies on gene expression (Zhu et al. 2019)(Buratin et al. 2020). Notably, unlike for FPs, experimental validations cannot amend the bias derived by FN errors. These considerations prompted us to consider a method evaluation metric that equally weighted precision and recall, such as the F_1_-score.

Interestingly, the evaluation by F_1_-score in our simulated data analysis highlighted that some methods with diverging precision and recall, such as CIRI2 and Segemehl (Figure 3a), were equally ranked. Moreover, the real data set analysis showed an ample recall variation of each method across the samples, while precision scores were tighter both between and within methods. These results confirmed that choosing a single method for circRNA discovery is problematic since no single best performing method exists. We showed that an approach leveraging the advantages brought by each of the integrated methods allows CirComPara2 to perform better than the single methods consistently in different experimental contexts. Besides, we showed that CirComPara2 predictions are more inclusive and robust than single method ones, even upon low count filtering, indicating that the recovered circRNAs missed by other methods are not merely of low abundance.

To our knowledge, CirComPara has been the first bioinformatics tool to combine multiple circRNA detection methods in an automated software pipeline. Since its former implementation, the continuous upgrade of CirComPara evolved into a substantially improved new tool, CirComPara2, with an efficacious method combination strategy. Other computational pipelines that employed multiple circRNA detection tools, such as RAISE (Li et al. 2017a) and circRNAwrap (Li et al. 2019), considered the circRNAs predicted by the union or the intersection of all methods, respectively, without validating which method combination was best. Our data show that those two approaches could be suboptimal as they suffer either low precision or sensitivity, whereas the best tradeoff between precision and recall is achieved with predictions of methods taken pairwise.

Recently, large circRNA databases have been compiled using method combination strategies (Vromman et al. 2020). For instance, circAtlas2.0 (Wu et al. 2020) retained circRNAs identified by at least two methods among CIRI2, CIRCexplorer2, Findcirc and DCC; the same combination but replacing DCC with circRNA_finder has been used in CircRic (Ruan et al. 2019). Applying these two approaches to our real-data sets analysis, we observed that they performed better than most of the algorithms except CirComPara2 (Supplementary Table S2), which achieved similar precision but slightly more than 0.2 better recall. Such a result suggests that CirComPara2 could be employed to compile comprehensive and reliable circRNA databases in future works.

Importantly, the computational pipelines used to compile circAtlas2.0 and CircRic were not implemented as software tools available to the scientific community. Instead, we made the automated and computationally efficient CirComPara2 pipeline ready-to-use and portable through a Docker container, freeing bioinformaticians from the several difficulties posed by implementing a computational pipeline, such as installation of multiple tools, software portability, code maintenance and documentation (Menegidio et al. 2018).

CirComPara2 is the only tool that aggregates various method expression estimates into unified values that eliminate redundant counts of BJR identified by multiple tools without relying on additional re-alignment of the reads. CircRNAwrap, RAISE, and CircAtlas2.0 applied a quantification step downstream of circRNA detection to estimate the circRNA expression. They considered the re-alignment of the reads onto pseudo reference circRNA sequences through Sailfish-cir (Li et al. 2017b), RAISE itself, and CIRIquant (Zhang et al. 2020), respectively, thus increasing the computational requirements of the pipeline. Nevertheless, both Sailfish-cir and CIRIquant can be easily applied downstream to CirComPara2 predictions.

CircRNA studies are likely to grow in several branches of biology, both on model (Weigelt et al. 2020) and non-model organisms and beyond the biomedical field (Wu et al. 2021a; Chu et al. 2021; Liang et al. 2019), prompting the development of improved tools allowing more extensive circRNA investigation to unravel circRNA-related condition peculiarities, such as differential circRNA expression (Gaffo et al. 2019; Wu et al. 2021c; Tian et al. 2020), imbalances of the CLP (Buratin et al. 2020), and prevailing circular transcript isoform expression (Izuogu et al. 2018). CirComPara2 is a resource tool meeting these needs, as already proved by the successful application of its embryonic implementations in several studies of human diseases and of other species, including plants (Gaffo et al. 2019; Frydrych Capelari et al. 2019; Buratin et al. 2020; Dal Molin et al. 2020).

In this work, we described the main features of CirComPara2, our automated and computationally efficient software pipeline for circRNA expression characterisation that also allows traditional gene expression analysis and the computation of circRNA to host-gene linear transcript abundance. Importantly, we demonstrated the asset given by the CirComPara2 method combination strategy for circRNA discovery, which provides robust and inclusive predictions in diverse biological contexts. With CirComPara2, we aim to provide a helpful bioinformatics tool to obtain a more comprehensive picture of transcriptomes and boost the understanding of circRNA features, biological and pathogenetic roles.

## Methods

### Simulated data set

CircRNA reads were simulated with the CIRI_simulator from the CIRI2 tool suite using the whole GRCh38 human genome and Gencode v29 gene annotation.

The parameters used in ciri_simulator were: -C 20 -LC 0 -R 1 -LR 1 -L 101 -E 1 -CHR1 0 -M 250 - M2 450 -PM 0 -S 70 -S2 0 -SE 0 -PSI 10.

A total of 10% (30% for the highly pruned annotation simulation) of the gene annotation for the simulated circRNA parent genes was removed from the annotation file and used as input to circRNA detection methods and to simulate linear transcript reads with Polyester (the reads_per_transcript parameter was set to 300). The linear transcript read files were then concatenated to the circRNA read files.

Code for generating the simulated data is available as a software tool *CCP_simulator* at https://github.com/egaffo/CCP_simulator. The parameter ANNOPARTS used in *CCP_simulator* for the two filters on gene annotation (standard and high pruning) was set to “85,4,5,1,0,0,0” and “60,15,10,2.5,6,3.5,3”, respectively.

### Real-data sets

Overall, 71 samples with matched rRNA depletion and rRNA depletion followed by RNase R treatment libraries from six studies (Table 1), for a total of 142 samples processed, were retrieved from Gene Expression Omnibus (GEO) or the National Genomics Data Center (NGDC) databases. Reads from PRJCA000751 were trimmed to 150bp in the preprocessing phase, as reported in the original work.

Method predictions in genomic scaffolds or from the mitochondrial genome were not considered. CircRNAs predicted with a length shorter than the library read length or longer than the longest gene expressed in the sample (computed as genes with TPM ≥ 1 computed by StringTie v2.1.4) were filtered out.

### CircRNA detection methods’ parameters

The following genome and gene annotations from the Ensembl database were used in the analyses: GRCh38 human genome and v97 gene annotation, Mmul 10 *Macaca mulatta* genome and v101 gene annotation, and GRCm38 *Mus musculus* genome and v101 gene annotation.

CirComPara2 default parameters were set for the analyses, which are as follows: adaptors from the Trimmomatic v0.39 TruSeq3-PE-2.fa file; PREPROCESSOR = “trimmomatic”; PREPROCESSOR_PARAMS = “MAXINFO:40:0.5 LEADING:20 TRAILING:20 SLIDINGWINDOW:4:30 MINLEN:50 AVGQUAL:26”. For the PRJCA000751 data sets, the

CROP:150 option was appended to the parameter. STAR_PARAMS = ‘--runRNGseed 123 -- outSJfilterOverhangMin 15 15 15 15 --alignSJoverhangMin 15 --alignSJDBoverhangMin 15 -- seedSearchStartLmax 30 --outFilterScoreMin 1 --outFilterMatchNmin 1 --outFilterMismatchNmax 2 --chimSegmentMin 15 --chimScoreMin 15 --chimScoreSeparation 10 --chimJunctionOverhangMin 15’. CIRCRNA_METHODS = ‘circexplorer2_bwa, circexplorer2_segemehl, circexplorer2_star, circexplorer2_tophat, ciri, dcc, findcirc’ (‘circrna_finder’ and ‘testrealign’ values were used in additional runs to obtain CircRNA_finder and Segemhel predictions); CPUS = 12; BWA_PARAMS = ‘-T 19’; SEGEMEHL_PARAMS = ‘-D 0’; BOWTIE2_PARAMS = ‘--reorder --score-min=C,-15,0 –q --seed 123’; DCC_EXTRA_PARAMS = ‘-fg -M -F -Nr 1 1 -N’; TESTREALIGN_PARAMS = ‘q median_1’; FINDCIRC_EXTRA_PARAMS = ‘--best-qual 40 --filter-tags UNAMBIGUOUS_BP -- filter-tags ANCHOR_UNIQUE’ (this setting implements the optimization suggested by Hansen (Hansen 2018)). MIN_METHODS = 2; MIN_READS = 2; CIRC_MAPPING = “{‘SE’: [‘STAR’,’TOPHAT’,’BOWTIE2’],’PE’:[‘BWA’,’SEGEMEHL’]}”; HISAT2_PARAMS = ‘--seed 123’.

Segemehl predictions reported in the sngl.bed and trns.txt files were merged to include spliced reads spanning >20,000 bps, as CirComPara2 performs it.

Read alignment and circRNA detection methods’ parameters were set as the value in the corresponding CirComPara2 parameters. All other parameters not mentioned were left with default values.

### Details of the CirComPara2 method

CircRNA expression estimates in CirComPara2 are represented as the sum of all unique back-splice junction reads (BJRs) identified by the circRNA methods. Then, the number of BJR fragments is counted while keeping track of the number of methods detecting each circRNAs. Finally, circRNAs with at least two reads identified by two or more methods are reported. The MIN_METHODS and MIN_READS options can be used to modify the required minimum reads and methods.

The CirComPara2 pipeline considers a preliminary alignment step that maps the reads linearly on the genome. Linearly aligned reads are then used to characterise canonical gene and transcript expression and count the linearly spliced reads spanning the back-splice junctions. Instead, linearly unmapped reads are used as input to the circRNA detection methods.

### Evaluation metrics and statistical tests

In evaluating method predictions, to compare samples with their matched control libraries, the different sequencing depths of the control library and the biochemical variability of the exoribonuclease in RNase R treatment have been adjusted by considering the proportion between the expression of the predicted circRNAs and the linear transcripts sharing the back-spliced exons. In each sample and for each method, we computed the circular-to-linear expression proportion (CLP) of each predicted circRNA by counting the number of reads back-spliced (BS_reads_) and linearly-spliced (LS_reads_) on the circRNA back-splice junctions, respectively. Then, the CLPs were calculated as BS_reads_/(BS_reads_+ LS_reads_). Szabo and Salzman (Szabo and Salzman 2016) also suggested using the ratio between the expression of circRNAs and their linear counterparts to overcome the evaluation issues deriving by RNase R treated control samples.

Assuming that RNase R should degrade linear transcripts more than circRNAs regardless of its efficiency in different samples, circRNAs with an equal or increased CLP in the control samples were deemed true-positives (TPs); false-positives (FPs) otherwise. Only circRNAs detected in the control libraries were considered to limit incorrect FP calls due to a lower sequencing depth of the control sample and to RNase R sensitive circRNAs (Jeck et al. 2013).

The precision was defined as TP/(TP + FP). The recall was computed as TP/(TP + FN) and F_1_-score as (2 × Precision × Recall)/(Precision + Recall), where TP and FN denote the true-positive and false-negative numbers.

Wilcoxon one-tailed paired tests were used to compare CirComPara2 greater recall and F _1_-score or lower precision with each method. Bonferroni multiple test correction was applied to compute adjusted p-values (q-values). The q-values reported in the main text refer to the highest value among the pairwise comparisons for recall and F_1_-score, whereas to the lowest value for precision comparisons.

### Software versions

The software version of the circRNA detection methods and chimeric read aligners used in this study: CIRCexplorer2 (Zhang et al. 2016) v2.3.8, CircRNA_finder (Westholm et al. 2014) v1.1, CIRI2 (Gao et al. 2018) v2.0.6, DCC v0.4.8 (Cheng et al. 2016), Findcirc (Memczak et al. 2013) v1.2, Segemehl (Hoffmann et al. 2014) v0.3.4 (used also by CIRCexplorer2), BWA MEM (Li 2013) v0.7.15 (used by CIRI2 and CIRCexplorer2), STAR (Dobin et al. 2013) v2.6.1e (used by circRNA_finder, DCC, and CIRCexplorer2), TopHat2 (Kim and Salzberg 2011) v2.1.0 (used for the TopHat-Fusion algorithm by CIRCexplorer2). CirComPara2 v0.1.2.1 was used to run the analysis.

## Data access

All the analysed data have been previously published and the accessions are referenced in Table 1.

## Acknowledgements

We thank Dr Geertruij te Kronnie for insightful suggestions and critical revision of the manuscript.

## Author Contributions

Conceptualisation, E.G.; Data curation, E.G.; Formal Analysis, E.G. and A.B.; Funding Acquisition, E.G. and S.B.; Methodology, E.G. and A.B.; Project Administration, S.B.; Resources, S.B.; Software, E.G. and A.B.; Supervision, E.G. and S.B.; Visualization, E.G.; Writing - Original Draft, E.G., A.B. and S.B.; Writing - Review & Editing, E.G., A.B., A.D.M. and S.B. All authors read and approved the final manuscript.

