## Supplemental figures and tables for "Sensitive, reliable, and robust circRNA detection from RNA-seq with CirComPara2"

### Supplemental Material

**Supplementary Figures** **1**
Supplementary Figure S1 1
Supplementary Figure S2 2
Supplementary Figure S3 3

**Supplementary Tables** **3**
Supplementary Table S1 4
Supplementary Table S2 7

### Supplementary Figures

*Supplementary Figure S1*

The percentage of false-positive predictions (FP) in common between pairs of methods and FPs of every single method (in the diagonal). Findcirc is not shown as it had no false-positive predictions. Shapes and colour-scale highlight percentage values: cyan circles denote low values, narrower ellipses correspond to higher percentages (blue, magenta, to red).

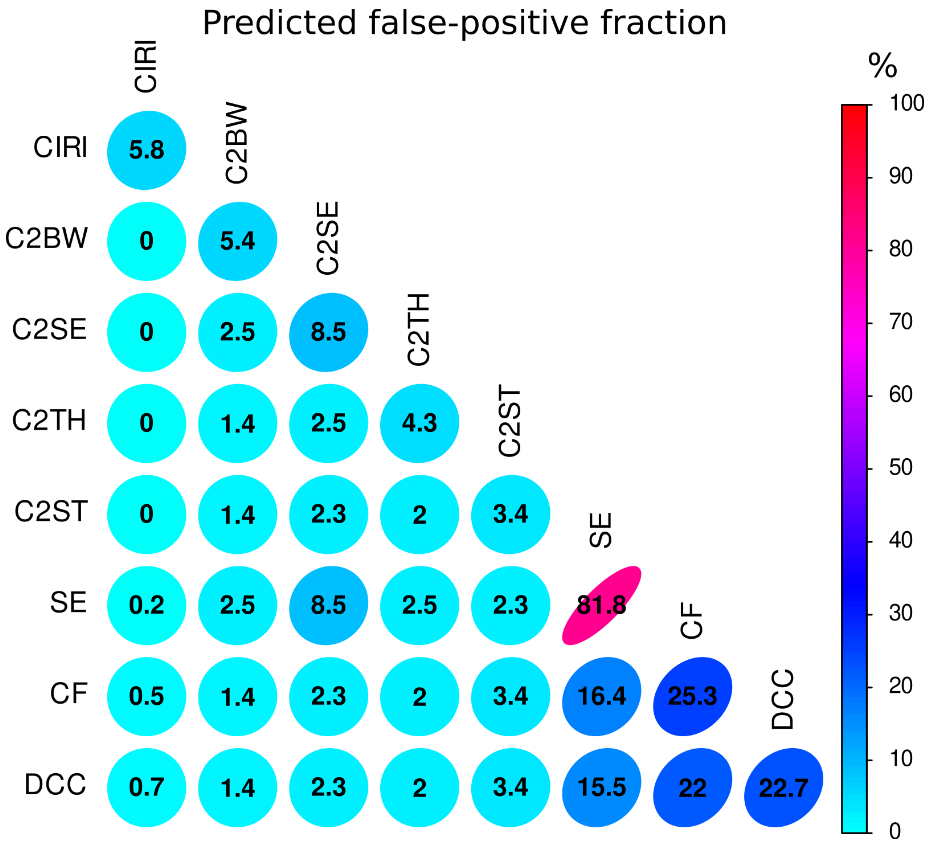

*Supplementary Figure S2*

Methods precision and recall calculated on the simulated data set using reduced gene annotation. Dashed curves delimit areas of $F_1$ score thresholds.

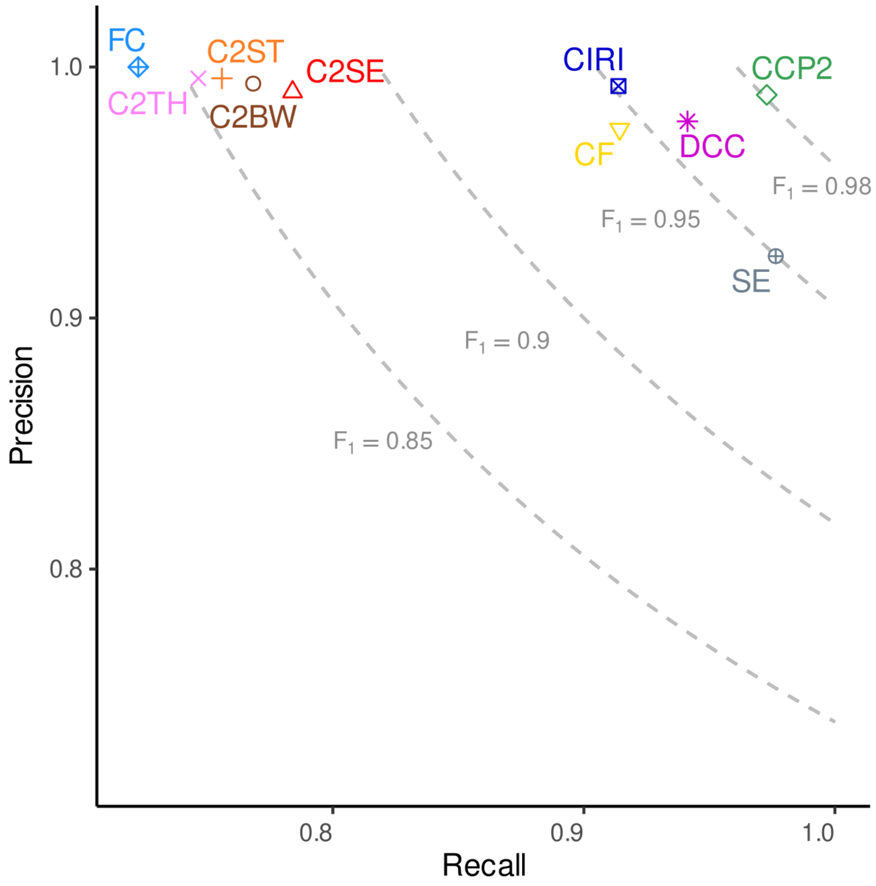

*Supplementary Figure S3*

Overall view of each method precision and recall for increasing thresholds of minimum back-splice junction read count filter (from 2 to 10, label in boxes).

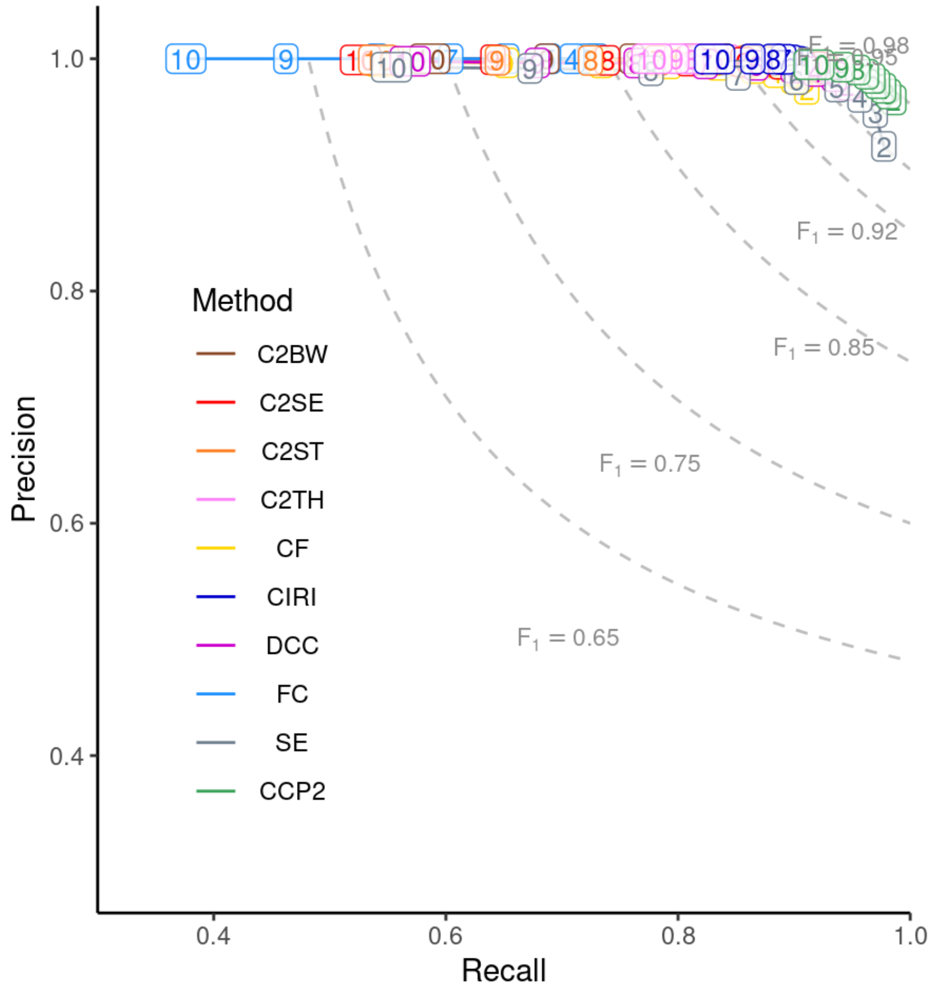

#### Supplementary Tables

*Supplementary Table S1*

Precision, recall and $F_1$ score of each circRNA detection method upon increasing thresholds (2 to 10) of minimum backsplice read count supporting the detection on the simulated data set. C2BW: CIRCexplorer2 on BWA; C2SE: CIRCexplorer2 on Segemehl; C2ST: CIRCexplorer2 on STAR; C2TH: CIRCexplorer2 on TopHat-Fusion; CCP2: CirComPara2; CF: circRNA_finder; CIRI: CIRI2; FC: Findcirc; SE: Segemehl

| Reads threshold | Method | Precision | Recall | $F_1$ score |
|---|---|---|---|---|
| ≥ 2 | *CCP2* | 0.99 | 0.98 | 0.99 |
| ≥ 2 | DCC | 0.98 | 0.95 | 0.96 |
| ≥ 2 | C2SE | 0.99 | 0.93 | 0.96 |
| ≥ 2 | SE | 0.92 | 0.98 | 0.95 |
| ≥ 2 | CIRI | 0.99 | 0.91 | 0.95 |
| ≥ 2 | C2BW | 0.99 | 0.90 | 0.95 |
| ≥ 2 | C2ST | 1.00 | 0.90 | 0.94 |
| ≥ 2 | CF | 0.97 | 0.91 | 0.94 |
| ≥ 2 | C2TH | 1.00 | 0.88 | 0.93 |
| ≥ 2 | FC | 1.00 | 0.73 | 0.84 |
| ≥ 3 | *CCP2* | 0.99 | 0.97 | 0.98 |
| ≥ 3 | SE | 0.95 | 0.97 | 0.96 |
| ≥ 3 | DCC | 0.98 | 0.93 | 0.96 |
| ≥ 3 | C2SE | 0.99 | 0.92 | 0.95 |
| ≥ 3 | CIRI | 0.99 | 0.91 | 0.95 |
| ≥ 3 | C2BW | 1.00 | 0.89 | 0.94 |
| ≥ 3 | CF | 0.98 | 0.90 | 0.94 |
| ≥ 3 | C2ST | 1.00 | 0.89 | 0.94 |
| ≥ 3 | C2TH | 1.00 | 0.87 | 0.93 |
| ≥ 3 | FC | 1.00 | 0.72 | 0.84 |
| ≥ 4 | *CCP2* | 0.99 | 0.97 | 0.98 |
| ≥ 4 | SE | 0.97 | 0.96 | 0.96 |
| ≥ 4 | DCC | 0.99 | 0.92 | 0.95 |
| ≥ 4 | CIRI | 1.00 | 0.91 | 0.95 |

| | | | | |
|---|---|---|---|---|
| ≥ 4 | C2SE | 0.99 | 0.91 | 0.95 |
| ≥ 4 | C2BW | 1.00 | 0.88 | 0.93 |
| ≥ 4 | CF | 0.99 | 0.88 | 0.93 |
| ≥ 4 | C2ST | 1.00 | 0.87 | 0.93 |
| ≥ 4 | C2TH | 1.00 | 0.85 | 0.92 |
| ≥ 4 | FC | 1.00 | 0.71 | 0.83 |

| | | | | |
|---|---|---|---|---|
| ≥ 5 | *CCP2* | 0.99 | 0.97 | 0.98 |
| ≥ 5 | SE | 0.98 | 0.94 | 0.96 |
| ≥ 5 | CIRI | 1.00 | 0.90 | 0.95 |
| ≥ 5 | DCC | 0.99 | 0.90 | 0.94 |
| ≥ 5 | C2SE | 1.00 | 0.89 | 0.94 |
| ≥ 5 | C2BW | 1.00 | 0.86 | 0.93 |
| ≥ 5 | CF | 0.99 | 0.87 | 0.92 |
| ≥ 5 | C2ST | 1.00 | 0.85 | 0.92 |
| ≥ 5 | C2TH | 1.00 | 0.84 | 0.91 |
| ≥ 5 | FC | 1.00 | 0.69 | 0.82 |

| | | | | |
|---|---|---|---|---|
| ≥ 6 | *CCP2* | 1.00 | 0.96 | 0.98 |
| ≥ 6 | CIRI | 1.00 | 0.90 | 0.94 |
| ≥ 6 | SE | 0.98 | 0.90 | 0.94 |
| ≥ 6 | DCC | 0.99 | 0.87 | 0.93 |
| ≥ 6 | C2SE | 1.00 | 0.86 | 0.92 |
| ≥ 6 | C2BW | 1.00 | 0.84 | 0.91 |
| ≥ 6 | C2TH | 1.00 | 0.83 | 0.91 |
| ≥ 6 | CF | 0.99 | 0.83 | 0.91 |
| ≥ 6 | C2ST | 1.00 | 0.82 | 0.90 |
| ≥ 6 | FC | 1.00 | 0.65 | 0.79 |

| | | | | |
|---|---|---|---|---|
| ≥ 7 | *CCP2* | 1.00 | 0.95 | 0.98 |
| ≥ 7 | CIRI | 1.00 | 0.89 | 0.94 |
| ≥ 7 | SE | 0.99 | 0.85 | 0.91 |
| ≥ 7 | C2TH | 1.00 | 0.82 | 0.90 |
| ≥ 7 | DCC | 1.00 | 0.82 | 0.90 |
| ≥ 7 | C2BW | 1.00 | 0.81 | 0.90 |
| ≥ 7 | C2SE | 1.00 | 0.81 | 0.89 |
| ≥ 7 | CF | 1.00 | 0.79 | 0.88 |

| | | | | |
|---|---|---|---|---|
| ≥ 7 | C2ST | 1.00 | 0.78 | 0.88 |
| ≥ 7 | FC | 1.00 | 0.60 | 0.75 |

| | | | | |
|---|---|---|---|---|
| ≥ 8 | *CCP2* | 1.00 | 0.95 | 0.97 |
| ≥ 8 | CIRI | 1.00 | 0.88 | 0.94 |
| ≥ 8 | C2TH | 1.00 | 0.81 | 0.90 |
| ≥ 8 | SE | 0.99 | 0.78 | 0.87 |
| ≥ 8 | DCC | 1.00 | 0.76 | 0.86 |
| ≥ 8 | C2BW | 1.00 | 0.76 | 0.86 |
| ≥ 8 | C2SE | 1.00 | 0.74 | 0.85 |
| ≥ 8 | CF | 1.00 | 0.73 | 0.85 |
| ≥ 8 | C2ST | 1.00 | 0.72 | 0.84 |
| ≥ 8 | FC | 1.00 | 0.54 | 0.70 |

| | | | | |
|---|---|---|---|---|
| ≥ 9 | *CCP2* | 1.00 | 0.93 | 0.96 |
| ≥ 9 | CIRI | 1.00 | 0.86 | 0.93 |
| ≥ 9 | C2TH | 1.00 | 0.80 | 0.89 |
| ≥ 9 | C2BW | 1.00 | 0.69 | 0.81 |
| ≥ 9 | DCC | 1.00 | 0.68 | 0.81 |
| ≥ 9 | SE | 0.99 | 0.67 | 0.80 |
| ≥ 9 | CF | 1.00 | 0.65 | 0.79 |
| ≥ 9 | C2ST | 1.00 | 0.64 | 0.78 |
| ≥ 9 | C2SE | 1.00 | 0.64 | 0.78 |
| ≥ 9 | FC | 1.00 | 0.46 | 0.63 |

| | | | | |
|---|---|---|---|---|
| ≥ 10 | *CCP2* | 1.00 | 0.91 | 0.95 |
| ≥ 10 | CIRI | 1.00 | 0.83 | 0.91 |
| ≥ 10 | C2TH | 1.00 | 0.78 | 0.87 |
| ≥ 10 | C2BW | 1.00 | 0.59 | 0.74 |
| ≥ 10 | DCC | 1.00 | 0.57 | 0.72 |
| ≥ 10 | SE | 0.99 | 0.55 | 0.71 |
| ≥ 10 | CF | 1.00 | 0.55 | 0.71 |
| ≥ 10 | C2ST | 1.00 | 0.54 | 0.70 |
| ≥ 10 | C2SE | 1.00 | 0.53 | 0.69 |
| ≥ 10 | FC | 1.00 | 0.38 | 0.55 |

*Supplementary Table S2*

**Supplementary Table S2.** Statistics of $F_1$-score, precision and recall achieved in the real data sets by each circRNA detection method. Methods are sorted according to descending median $F_1$-score. Qt: quartile; SEr: standard error; IQR: interquartile range.

| Method | $F_1$ Min | $F_1$ 1st Qt | $F_1$ Mean | $F_1$ SEr | $F_1$ Median | $F_1$ 3rd Qt | $F_1$ Max | $F_1$ IQR |
|---|---|---|---|---|---|---|---|---|
| CCP2 | 0.264 | 0.817 | 0.818 | 0.022 | 0.906 | 0.929 | 0.964 | 0.112 |
| SE | 0.335 | 0.634 | 0.740 | 0.017 | 0.822 | 0.861 | 0.915 | 0.228 |
| CircAtlas2 | 0.144 | 0.587 | 0.692 | 0.023 | 0.781 | 0.848 | 0.904 | 0.261 |
| CircRic | 0.144 | 0.585 | 0.691 | 0.023 | 0.778 | 0.847 | 0.904 | 0.262 |
| C2BW | 0.202 | 0.571 | 0.681 | 0.021 | 0.757 | 0.823 | 0.877 | 0.252 |
| CIRI2 | 0.168 | 0.501 | 0.666 | 0.025 | 0.741 | 0.854 | 0.930 | 0.352 |
| CF | 0.182 | 0.534 | 0.643 | 0.020 | 0.714 | 0.793 | 0.839 | 0.259 |
| C2SE | 0.182 | 0.558 | 0.648 | 0.019 | 0.704 | 0.777 | 0.869 | 0.218 |
| DCC | 0.182 | 0.511 | 0.633 | 0.020 | 0.702 | 0.771 | 0.842 | 0.260 |
| C2TH | 0.159 | 0.524 | 0.623 | 0.020 | 0.685 | 0.743 | 0.844 | 0.219 |
| C2ST | 0.159 | 0.483 | 0.596 | 0.019 | 0.655 | 0.726 | 0.785 | 0.242 |
| FC | 0.105 | 0.274 | 0.376 | 0.015 | 0.390 | 0.477 | 0.608 | 0.204 |

| Method | Recall Min | Recall 1st Qt | Recall Mean | Recall SEr | Recall Median | Recall 3rd Qt | Recall Max | Recall IQR |
|---|---|---|---|---|---|---|---|---|
| CCP2 | 0.304 | 0.752 | 0.821 | 0.016 | 0.860 | 0.922 | 0.963 | 0.170 |
| SE | 0.349 | 0.660 | 0.703 | 0.014 | 0.754 | 0.778 | 0.885 | 0.118 |
| CircAtlas2 | 0.143 | 0.478 | 0.616 | 0.020 | 0.650 | 0.744 | 0.850 | 0.266 |
| CircRic | 0.143 | 0.475 | 0.615 | 0.020 | 0.648 | 0.742 | 0.850 | 0.266 |
| C2BW | 0.205 | 0.474 | 0.597 | 0.017 | 0.626 | 0.707 | 0.798 | 0.233 |
| CIRI2 | 0.159 | 0.446 | 0.592 | 0.024 | 0.630 | 0.753 | 0.899 | 0.307 |
| CF | 0.171 | 0.428 | 0.543 | 0.017 | 0.578 | 0.664 | 0.732 | 0.236 |
| C2SE | 0.177 | 0.455 | 0.548 | 0.015 | 0.569 | 0.646 | 0.785 | 0.191 |
| DCC | 0.167 | 0.424 | 0.529 | 0.016 | 0.564 | 0.637 | 0.736 | 0.213 |
| C2TH | 0.140 | 0.431 | 0.510 | 0.015 | 0.526 | 0.609 | 0.745 | 0.178 |
| C2ST | 0.140 | 0.381 | 0.479 | 0.015 | 0.512 | 0.577 | 0.652 | 0.195 |
| FC | 0.072 | 0.186 | 0.249 | 0.010 | 0.246 | 0.322 | 0.440 | 0.136 |

| Method | Precision Min | Precision 1st Qt | Precision Mean | Precision SEr | Precision Median | Precision 3rd Qt | Precision Max | Precision IQR |
|---|---|---|---|---|---|---|---|---|
| CCP2 | 0.221 | 0.843 | 0.833 | 0.028 | 0.959 | 0.985 | 0.994 | 0.141 |
| SE | 0.230 | 0.777 | 0.824 | 0.025 | 0.923 | 0.969 | 0.990 | 0.192 |
| CircAtlas2 | 0.144 | 0.817 | 0.832 | 0.030 | 0.963 | 0.987 | 0.998 | 0.170 |
| CircRic | 0.144 | 0.817 | 0.832 | 0.030 | 0.963 | 0.987 | 0.998 | 0.170 |
| C2BW | 0.191 | 0.821 | 0.833 | 0.029 | 0.968 | 0.986 | 0.995 | 0.165 |
| CIRI2 | 0.171 | 0.877 | 0.834 | 0.030 | 0.968 | 0.985 | 0.997 | 0.109 |
| CF | 0.186 | 0.838 | 0.829 | 0.029 | 0.958 | 0.980 | 0.996 | 0.142 |
| C2SE | 0.188 | 0.821 | 0.834 | 0.029 | 0.963 | 0.986 | 0.996 | 0.165 |
| DCC | 0.183 | 0.838 | 0.830 | 0.029 | 0.957 | 0.983 | 0.997 | 0.145 |
| C2TH | 0.181 | 0.872 | 0.833 | 0.030 | 0.967 | 0.987 | 0.998 | 0.115 |
| C2ST | 0.183 | 0.862 | 0.836 | 0.029 | 0.964 | 0.987 | 0.998 | 0.125 |
| FC | 0.167 | 0.848 | 0.824 | 0.031 | 0.961 | 0.983 | 0.997 | 0.136 |
